# Deletion of exons 45 to 55 in the *DMD* gene: from the therapeutic perspective to the *in vitro* model

**DOI:** 10.1101/2023.09.13.557649

**Authors:** Javier Poyatos-García, Patricia Soblechero-Martín, Alessandro Liquori, Andrea López-Martínez, Elisa González-Romero, Rafael P. Vázquez-Manrique, Nuria Muelas, Gema García-García, Jessica Ohana, Virginia Arechavala-Gomeza, Juan J. Vílchez

## Abstract

Gene editing therapies in development for correcting out-of-frame *DMD* mutations in Duchenne muscular dystrophy aim to replicate benign spontaneous deletions. Deletion of 45**–**55 *DMD* exons (del45**–**55) was described in asymptomatic subjects, but recently serious skeletal and cardiac complications have been reported. Uncovering why a single mutation like del45–55 is able to induce diverse phenotypes and grades of severity may impact the strategies of emerging therapies. Cellular models are essential for this purpose, but their availability is compromised by scarce muscle biopsies. Here, we have introduced through CRISPR-Cas9 edition, a del45**–**55 mimicking the intronic breakpoints harboured by a subset of patients of this form of dystrophinopathy, into a Duchenne patient’s cell line. Dystrophin expression was restored in edited myoblasts and the myogenic defects were ameliorated. Besides confirming the potential of CRISPR-Cas9 to create tailored mutations as a useful approach to generate *in vitro* models, we also generated an immortalized myoblast line derived from a patient with a specific del45**–**55. Overall, we provide helpful resources to deepen into unknown factors responsible for DMD-pathophysiology.

**SUMMARY STATEMENT:** We restored dystrophin expression in a DMD culture by replicating the exact deletion in exons 45-55 harboured by mild patients, testing this therapeutic approach, and creating a new cell model.

## INTRODUCTION

Dystrophinopathies encompass a series of muscular disorders caused by mutations in the *DMD* gene causing structural or functional alterations in the protein dystrophin, a member of the dystrophin-glycoprotein complex (DGC) located in the sarcolemma that connects actin cytoskeleton to the extracellular matrix, conferring structural stability to the muscle fibres during contraction and having an important role in cellular signalling (Davies and Nowak, 2006; Rando, 2001). The variety of the phenotypic presentation of these disorders underlies the implication of different pathomechanisms, whose understanding is critical to focus medical management and treatment approaches for this subgroup of degenerative diseases.

The most severe and frequent phenotype consists in Duchenne muscular dystrophy (DMD), mainly produced by frame-disrupting *DMD* mutations. It is characterized by progressive skeletal muscle degeneration from early childhood, leading to loss of ambulation around 13 years of age and early death in the thirties due to respiratory and cardiac complications. On the other hand, mutations that preserve the *DMD* open reading frame (ORF) enable the production of partially functional dystrophins, usually causing Becker muscular dystrophy (BMD), defined by the preservation of ambulation beyond 16 years and a broader spectrum of the age of onset and severity of muscle weakness (Darras et al., 2022). Other phenotypes associated with *DMD* mutations include isolated hiperCKemia (Ferreiro et al., 2009), the presence of pseudo-metabolic manifestations like exercise-induced myalgia and rhabdomyolysis (Sanchez-Arjona, 2005), isolated cardiomyopathy (Nakamura, 2015), and cognitive and neurodevelopmental abnormalities usually associated to any of the aforementioned phenotypes (North et al., 1996). In addition to the impact of each mutation on the ORF, the clinical severity also depends on the alteration of the conformational structure of the resultant dystrophin that can impair its proper assembly and interaction with other proteins (Nicolas et al., 2015); and the influence of *trans* gene modifying factors (Bello and Pegoraro, 2019).

Currently, there is no effective cure for DMD but exon skipping and microdystrophin gene transfer are promising therapeutic approaches based on the restoration or replacement of the dystrophin functionality. Exon skipping can be achieved by restoring the reading frame using splice-switching antisense oligonucleotides (AONs) acting on pre-mRNA to promote the skipping of the targeted exon (Arechavala-Gomeza et al., 2012), or by gene editing technologies acting on the DNA producing a permanent excision of the exon/s of interest (Amoasii et al., 2018; Long et al., 2016; Nelson et al., 2016; Tabebordbar et al., 2016). On the other hand, genomic technologies to assemble crucial functional protein domains into constructs suitable to be borne by viral vectors are used for the microdystrophin gene transfer therapy (Duan, 2018). Nevertheless, although all these approaches have shown encouraging clinical and preclinical results, it is still essential to overcome important hurdles (such as specificity, efficiency, immunogenicity and delivery problems) in order to turn these approaches into real therapeutical options (Hammond et al., 2021; Happi Mbakam et al., 2022; Verhaart and Aartsma-Rus, 2019). In this regard, the analysis of BMD patients or other benign dystrophinopathy phenotypes with in-frame deletions mimicking those achieved by exon skipping is essential to predict the therapeutic potential of these approaches (Anthony et al., 2011a; Anthony et al., 2014a).

Among the different mutations along the *DMD* gene, the deletion of exons 45**–**55 (del45**–**55) has been postulated as an attractive therapeutic model capable to correct up to 47% of total DMD causing mutations, (Béroud et al., 2007; Flanigan et al., 2009; Nakamura et al., 2017). This in-frame deletion in the central dystrophin rod domain, formerly reported as benign, has been also shown to be causative of significant functional impairment, severe cardiac complications, cognitive alterations, and potential shortening of life expectancy (Poyatos-García et al., 2022). However, despite an extensive search, looking for potential *DMD cis* alterations (including intronic breakpoint positions disrupting regulatory sequences located in introns 44 and 55) and effects of *trans* modifying factors, the underlaying pathomechanisms still remain unrevealed (Poyatos-García et al., 2022). Thus, although functional dystrophin has successfully been restored in cellular and animal DMD-models through the skipping of exons 45**–**55, using both cocktails of AONs (Aoki et al., 2012; Echigoya et al., 2015; Echigoya et al., 2019; Lee et al., 2018) and CRISPR-Cas9 gene editing (Ousterout et al., 2015; Young et al., 2016; Young et al., 2017), it is still necessary to continue the investigation of this deletion using different approaches, to make this model an effective therapeutical alternative.

In this regard, cellular models are crucial, not only to investigate unidentified molecular mechanisms that contribute to the physiopathology, but also to develop new therapeutic strategies in preclinical steps. Primary human myoblasts isolated from patient muscle biopsies are the most reliable cellular models as they present the natural genomic background of the disease; however, they have limited proliferative capacity associated to cellular senescence. To overcome these issues, the immortalization of these myogenic human cell lines have demonstrated to be an important tool for neuromuscular disorders research, as they present increased proliferative capacity but maintain their differentiation potential and the myogenic expression pattern of the primary cells (Mamchaoui et al., 2011; Thorley et al., 2016). Nevertheless, in many hereditary neuromuscular disorders, muscle biopsies are not performed for diagnostic purposes due to the advance of the NGS-methods and consequently, the cell lines available of a given disease are limited, particularly those of a specific mutation. Thus, in order to solve this problem, the CRISPR-Cas9 system has emerged as a useful approach to create custom cell disease cell lines, and increasing the models available for the research of human neuromuscular diseases (Soblechero- Martín et al., 2021).

In this report, we evaluated, as a proof of concept, the replication of the intronic breakpoint positions of a subgroup of del45**–**55 patients previously analysed (Poyatos-García et al., 2022), using the CRISPR-Cas9 gene editing tool, as an attractive approach to restore the *DMD* reading frame of a DMD cell line. This strategy would also be useful to generate cell models with specific deletions that would help in deepen into novel disease pathomechanisms. In addition, we characterized an immortalized cell line derived from an asymptomatic patient harbouring that specific del45**–**55.

## RESULTS

### Generation of del45–55-D1 immortalized cell model (Im**Δ**45–55-D1)

Manipulating primary human myoblasts is challenging due to its low proliferative potential, so we decided to immortalize the primary cells from a 32-year-old male carrier with a del45–55 with specific intronic breakpoints. This patient was classified as asymptomatic as, though presenting raised serum creatin-kinase (CK) levels, no signs of functional impairment were observed. We named this specific deletion of *DMD* exons 45 to 55 as “del45–55-D1”, in accordance to results previously obtained by our group (Poyatos-García et al., 2022) and the culture derived from it “ImΔ45**–**55-D1”. Once immortalised, the proliferation potential of the primary and immortalized cell lines was analysed at 24, 48, 72, 96 h; confirming that the immortalized line proliferated significantly more than the primary one (Fig. S1).

### Generation of edited myoblasts clones mimicking the specific del45–55-D1 (Edited**Δ**45–55)

We replicated del45–55-D1 to restore the dystrophin expression in an immortalised DMD cell line with a deletion of exon 52 (DMDΔ52) using a CRISPR-Cas9 gene editing method. We compared this culture, that we named “EditedΔ45**–**55” with the immortalized myoblasts with the same mutation (ImΔ45**–**55-D1) to prove the suitability of this approach to generate cell models.

To achieve this purpose, we designed two CRISPR gRNA targeting each intronic breakpoint that were cloned into plasmid vectors expressing the Cas9 nuclease and GFP (Table S1). The four plasmids containing each gRNA, were transfected individually into HEK293 cells and their cleavage efficiencies were evaluated using the T7E1 assay. According to the estimated indel frequencies (Fig. S2), we chose sequences gRNA_44.1 and gRNA_55.2 (targeting intron 44 and 55 breakpoints, respectively) to generate the edited clones. Both plasmids were co-transfected into HEK293 cells confirming the production of the del45–55-D1. The location of the deletion across the introns 44 and 55, and the sequences and design of the gRNAs are represented in Fig. 1A,B.

**Figure 1.**
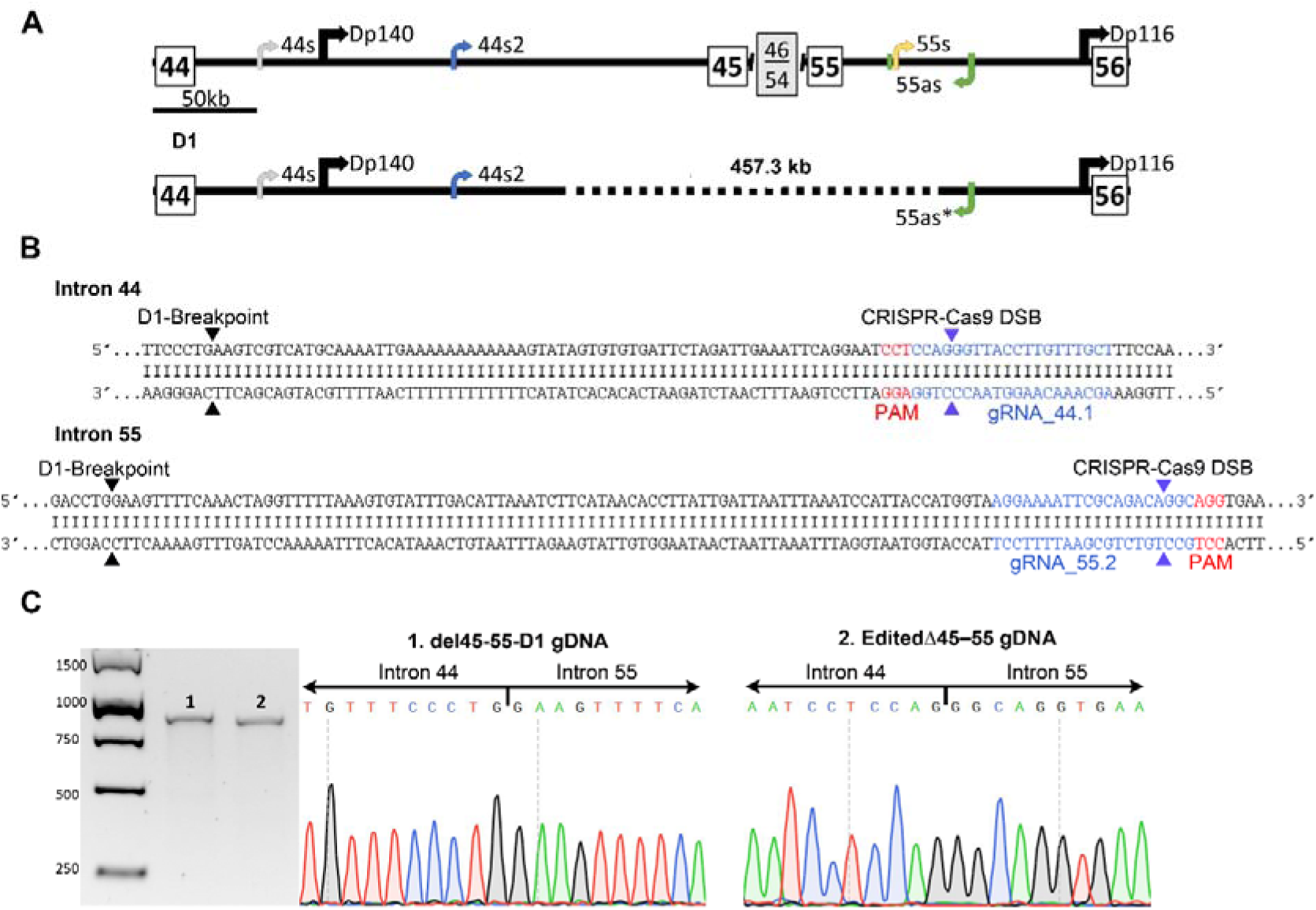
CRISPR-Cas9 design based on del45–55-D1 breakpoint location. (A) In the upper part is represented the genomic architecture along introns 44 and 55, showing the relative location of the promoter of the Dp140 and Dp116 dystrophin isoforms; as well as the position of the lncRNA 44s, 44s2, 55s and 55as. Below, there is a representation of the specific breakpoints in introns 44 and 55 of the deletion group D1 (del45–55-D1); and its implication over the commented elements (dotted lines indicate the deleted region). (B) Genomic sequences of introns 44 and 55, where the location of the patient breakpoints is indicated as black arrowheads. The gRNAs targeting each intron are represented blue letters, and its PAM (protospacer adjacent motif) sequences in red ones. Purple arrowheads indicate the location of the expected Cas9 DSB (double strand break). (C) Genomic sequences of the deletion junctions, confirmed by Sanger sequencing of (1) the patient harbouring this specific deletion and (2) the EditedΔ45–55 clone.

The plasmids containing the selected gRNAs were then co-transfected into the target immortalized human DMDΔ52 myoblasts. Forty-eight hours after transfection, GFP positive cells indicating that they have incorporated at least one of the vectors (14.42% of the total population) were sorted by FACS and individually seeded into 96-well plates; in order to obtain homogenous clonal cell cultures. The clones were expanded and screened by PCR and Sanger sequencing, identifying approximately one out ten with the specific del45–55-D1 (Fig. 1C). One positive clone was expanded for further characterisation.

### Cleavage efficiency and off-target events analysis

Amplicon deep sequencing analysis was performed to quantify the cleavage efficiencies (on-targets) of the selected gRNAs (gRNA_44.1 and gRNA_55.2) and to evaluate if they have produced any unspecific cleavage (off-targets events). We used the Cas-OFFinder and Breaking-Cas predictors (Bae et al., 2014; Oliveros et al., 2016) to identify the potential off-targets associated to each gRNA (Table 1).

**Table 1.**
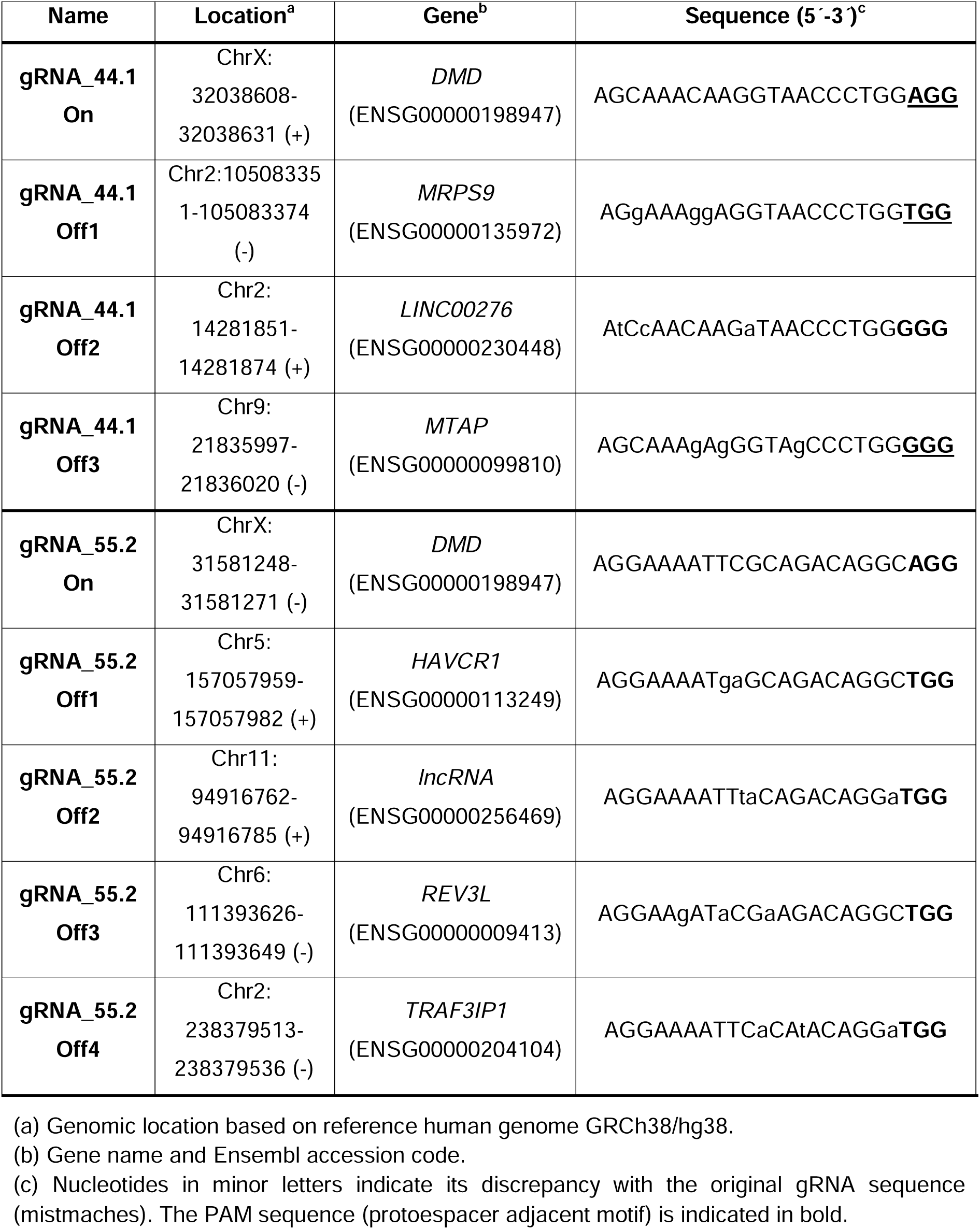
Sequence and genomic location of the on-target and potential off-targets regions.

Extracted DNA from transfected and untrasnfected cultures was used to generate the amplicon sequencing libraries. The fastaq.gz files from the Miseq sequencer (Illumina) were analysed with the CRISPResso2 software (Clement et al., 2019) that uses a narrow window, restricted to the expected cleavage site, to quantify the DNA modifications (insertions, deletions and substitutions).

All amplicons (from both transfected and untransfected cells) showed high coverage (above 400,000 properly aligned reads) except for gRNA_44.2 off-target 3 amplicon, which did not meet the software coverage requirements for its analysis. Probably, it might have occurred due to an error in the concentration determination of this amplicon, affecting in the final pool. The Table S2 shows the number of reads classified according to each DNA modification for each analysed amplicon.

Regarding the on-target, we observed a 5.50% and a 9.99% of modified reads associated to the gRNA_44.1 and 55.2, respectively. In both, the majority of modifications corresponded to indels, characteristic of the NHEJ repair pathway (Fig. 2).

**Figure 2.**
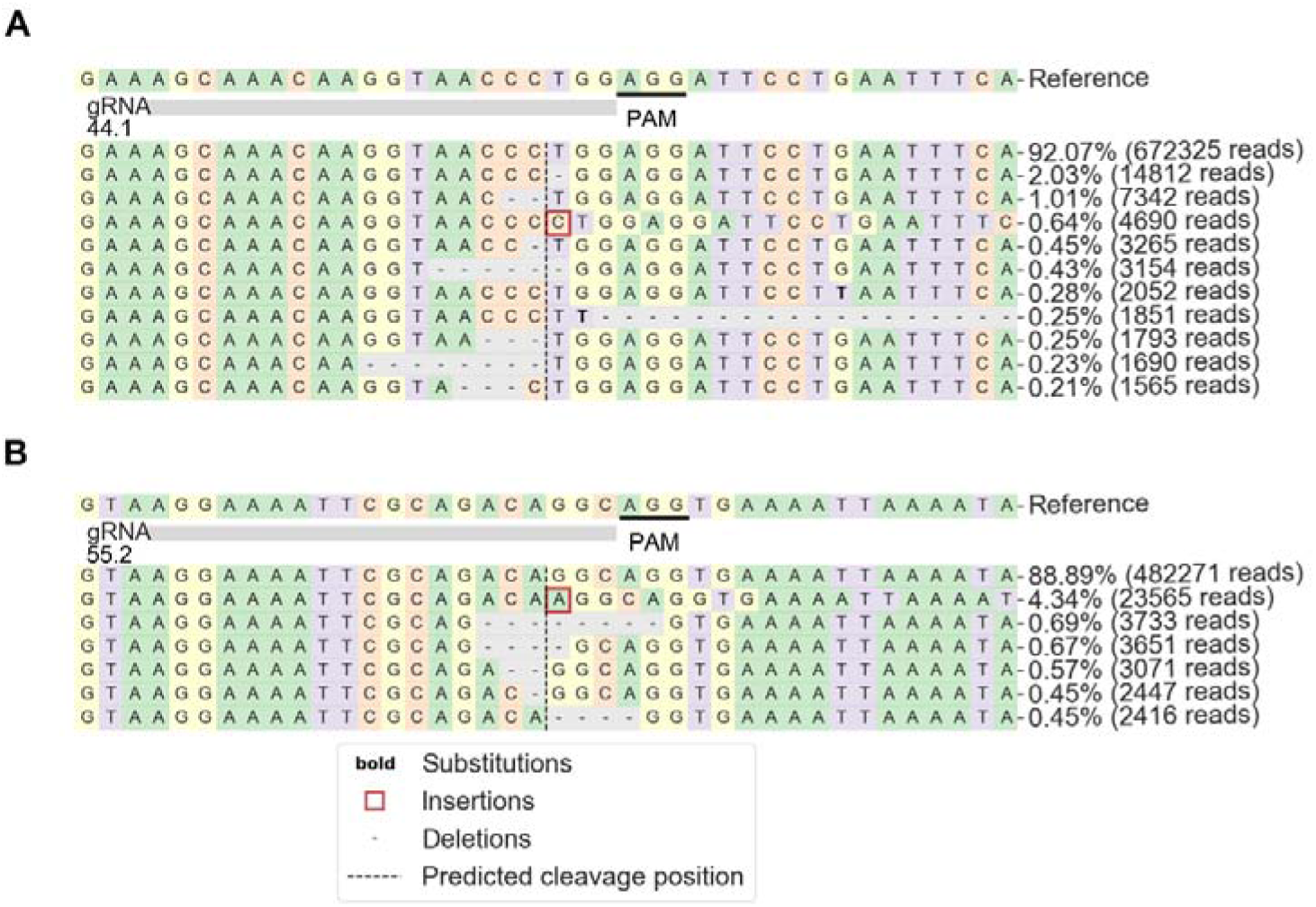
Summary of the principal allele frequencies generated after the gRNAs cleavages. Representation of the NGS reads frequencies derived from the transfection of gRNA44.1 (A) and gRNA55.2 (B). The reference sequence is indicated along with the principal modifications, where percentage is indicated. The image is adapted from the results obtained with the CRIPSPresso2 program. Figure legend explains the produced changes and the expected Cas9 cleavage site.

On the other hand, we used the CRISPResso2 Compare software to study the off- target events; as it analyses the results of the transfected and the untransfected sample with regard of the reference sequence for each amplicon (Fig. S3). We did not find any indel event near the expected cleavage site in any transfected sample. However, we did observe the presence of substitutions, that in some cases (gRNA_44.1 off-target 1 and gRNA_55.2 off-target 4) were enriched in the untransfected sample. We observed the presence of substitution on the transfected sample of gRNA_44.1 off-target 2; but they appeared very distant from the expected cleavage site (Fig. S3).

In addition, the seven potential off-targets were also analysed in the EditedΔ45–55 clone used for the functional characterization assays, through PCR followed by Sanger sequencing. No off-target effect was found at any locus analysed (Fig. S4).

### Dystrophin expression is recovered in the Edited**Δ**45–55 clone

We evaluated if the generation of the specific del45**–**55-D1 is able to restore the *DMD* reading frame of a DMDΔ52 cell line, enabling dystrophin production. In addition, we consider a valuable approach to compare the edited cell line to the immortalized myoblasts from the patient harbouring this specific deletion (ImΔ45**–**55-D1). For these experiments, we also introduced the original immortalized cell lines DMDΔ52 and control (C1).

First, RT-PCR Sanger sequencing of EditedΔ45–55 differentiated myotubes cDNA confirmed the in-frame deletion of 11 exons at RNA level (comparable to the template ImΔ45**–**55-D1sequence) (Fig. 3A). Additionally, the quantification of dystrophin cDNA confirmed the rescue of dystrophin expression in the EditedΔ45–55 clone, compared to the DMD unedited line, and presenting even greater amounts than control and ImΔ45**–** 55-D1 lines (Fig. 3B).

**Figure 3.**
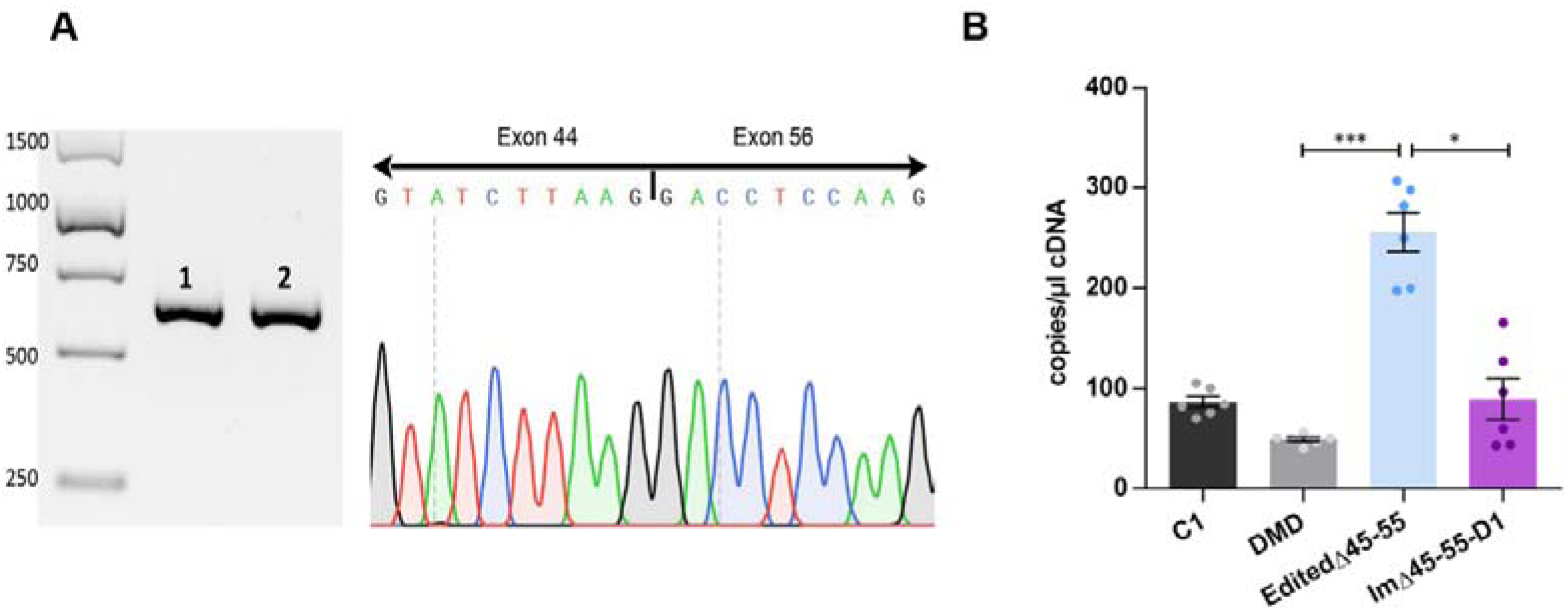
Dystrophin cDNA expression analysis of the edited clone. (A) Confirmation of the del45–55-D1 of the differentiated myotubes cDNA from (1) the ImΔ45–55-D1 and (2) from the EditedΔ45–55 clone. In both samples the exon 56 sequence is juxtaposed to the exon 44 one. (B) Dystrophin expression quantification through ddPCR of the differentiated myotubes cDNA from Control (C1), unedited DMD, the EditedΔ45–55 and the ImΔ45–55-D1 cell lines (n=8 replicates). The bar graphs represent mean ± SEM. *p < 0.05, ***p < 0.001, according to Kruskal-Wallis test.

Then, we proceeded to the dystrophin characterization at the protein level. The immunofluorescence of the EditedΔ45–55 differentiated myotubes confirmed the restoration of dystrophin expression, and its proper localization (Fig. 4A). We also performed a precise dystrophin quantification using in-cell western, confirming the dystrophin expression increase in the EditedΔ45–55 myotubes respect the non-edited DMD one’s (Fig. 4B).

**Figure 4.**
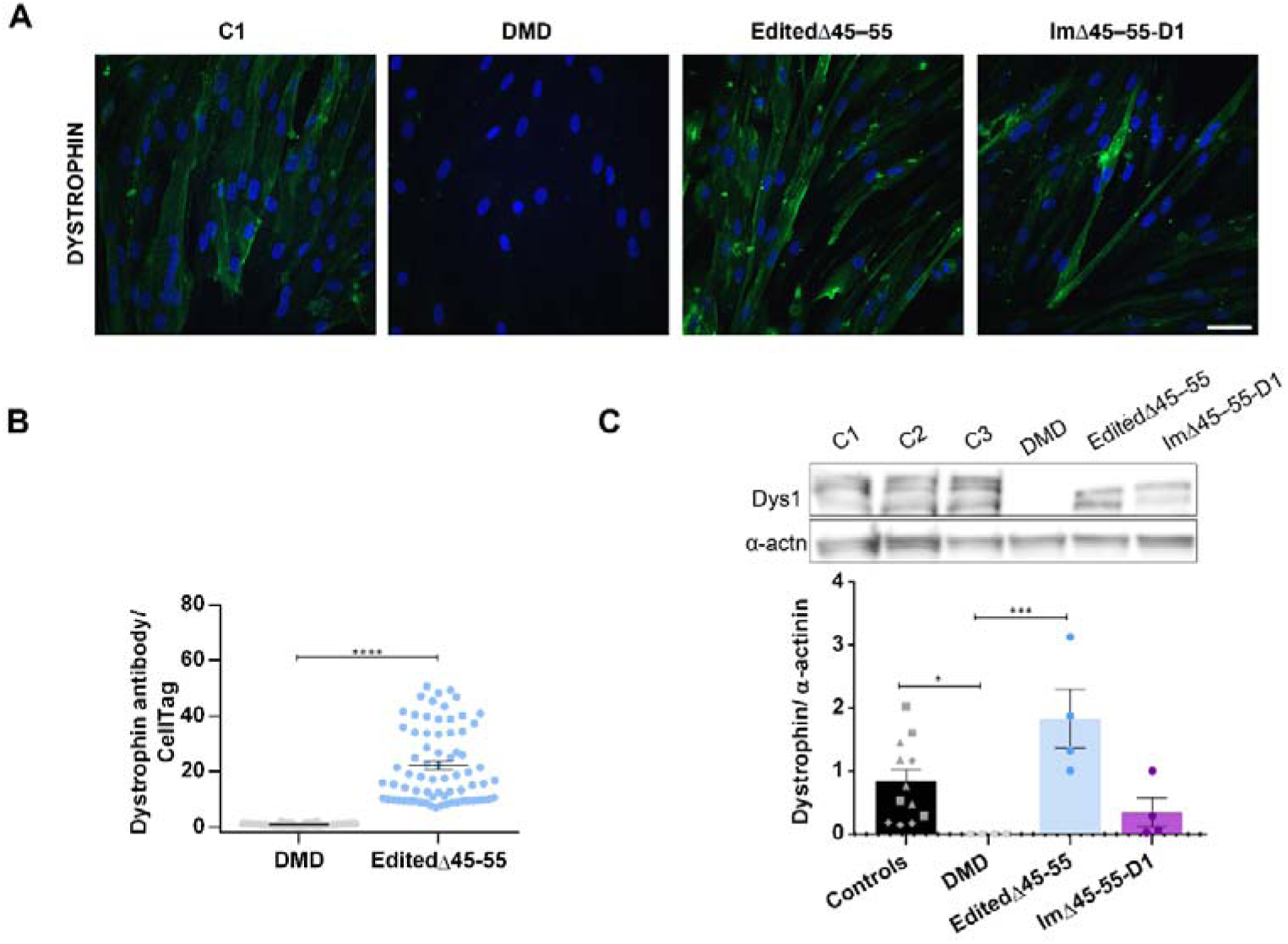
Dystrophin evaluation of the edited clone myotubes. **(A)** Representative confocal images of dystrophin immunofluorescence (green) in differentiated myotubes from: Control (C1), DMD, EditedΔ45–55 and ImΔ45–55-D1. Nuclei were stained with DAPI (blue) (scale bar=50µm). **(B)** In-cell western quantification of dystrophin expression of the EditedΔ45–55 myotubes compared to the unedited DMD ones. Dystrophin signal is normalized to cell number signal and set to 1 (Cell Tag) (n=72 wells). **(C)** Quantification and representative blots of dystrophin in protein extracts from 3 healthy controls (C1=, C2=, C3=); DMD, the EditedΔ45–55 and ImΔ45–55-D1 myotubes. Dystrophin levels (Dys1) were normalized to α-actinin signal (α-actn) (n=4 technical replicates). *p < 0.05, ***p < 0.001, ****p < 0.0001 according to Mann– Whitney U (B) and Kruskal-Wallis (C) and error bars represent mean ± SEM.

The assessment of dystrophin quantification was also carried out by western blot in the 4 mentioned cell lines, and two additional control samples (C2 and C3) were included. The dystrophin recuperation in the edited myotubes was confirmed, showing higher levels than the ImΔ45**–**55-D1 and the 3 controls myotubes’ (grouped) (Fig. 4C).

### The introduction of the del45–55-D1 by CRISPR-Cas9 rescues the myoblasts differentiation defects of DMD cells

Myoblasts derived from DMD patients harbouring out-of-frame mutations preventing dystrophin production, show important defects in their differentiation capability (Choi et al., 2021; Soblechero-Martín et al., 2021).

In order to evaluate this process, we analysed the myoblasts’ fusion capacity by immunofluorescence of Desmin, a muscle-specific type III intermediate filament essential for the proper muscular structure and function. We observed that, after 7 days differentiating, the DMD myotubes’ fusion index was significatively reduced compared to healthy control, while in the EditedΔ45–55 myotubes this parameter was recovered. We did not find any evidence between the patient ImΔ45**–**55-D1and control myotubes (Fig. 5A,B).

**Figure 5.**
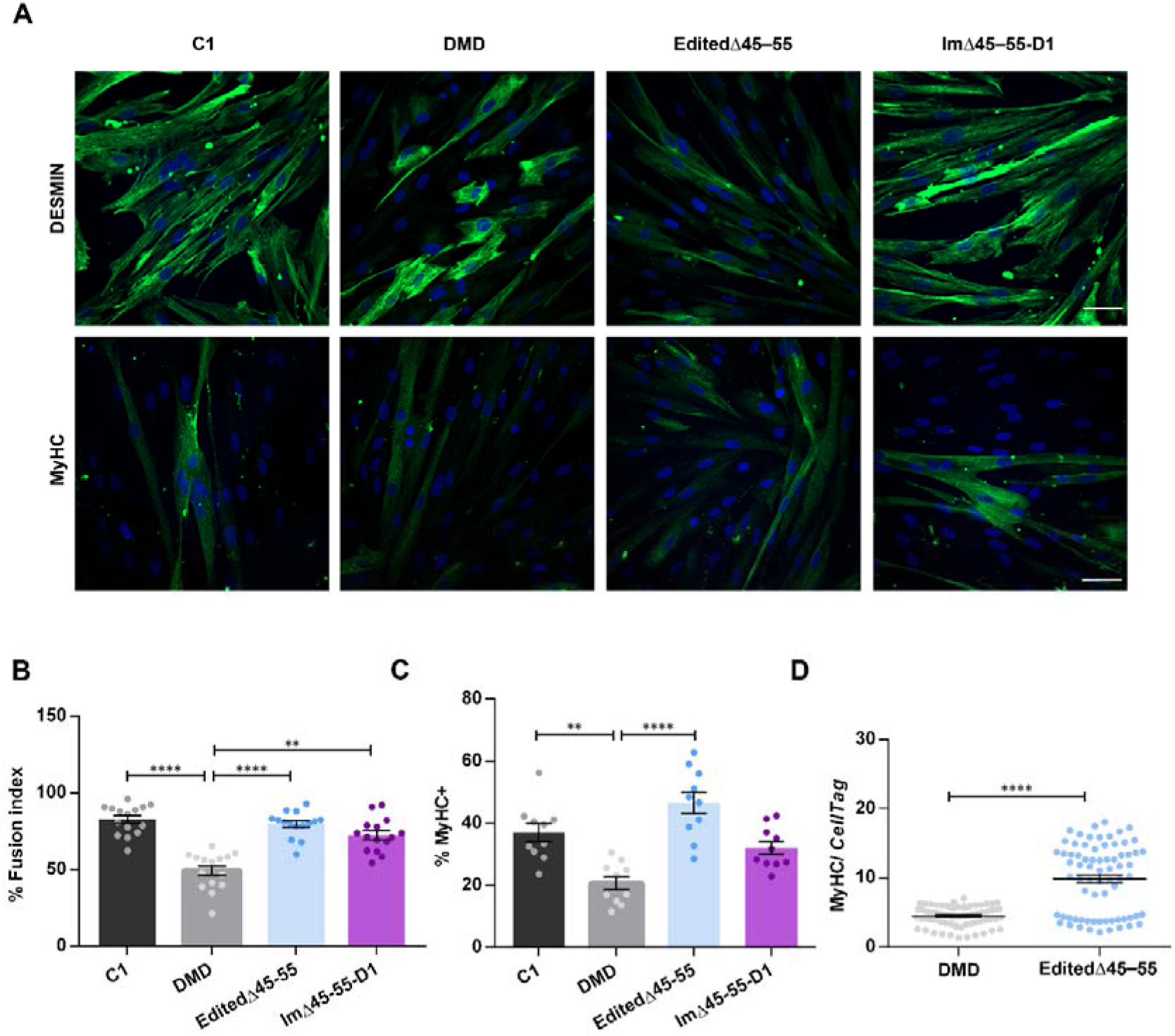
Evaluation of the myogenic differentiation process after gene edition. **(A)** Representative confocal images of Desmin and MyHC (myosin heavy chain) immunofluorescence (green) in control (C1), DMD, EditedΔ45–55 and ImΔ45–55-D1 differentiated myotubes. Nuclei were stained with DAPI (blue) (scale bar=50µm). **(B)** Quantification of the myogenic fusion index of the cell lines was calculated using the Desmin micrographs (15 images, 400-700 total nuclei). **(C)** Quantification of the percentage of nuclei within MyHC+ cells in each condition (10 images, 300-500 total nuclei). **(D)** In-cell western MyHC quantification of the EditedΔ45–55 myotubes compared to the unedited DMD ones. The bar graphs show the mean ± SEM; **p <0.01, ****p < 0.0001, according to Mann–Whitney U (B) and Kruskal-Wallis (C) tests.

Additionally, the myosin heavy chain (MyHC) immunofluorescence was performed as late-differentiation marker (Chal and Pourquié, 2017) in the differentiated myotubes of the 4 cell lines. We confirmed a differentiation defect in the DMD myotubes, as only 20% of the nuclei corresponded to MyHC+ cells, compared to the 37% of the controls’ nuclei. Similar to fusion index, this phenotype was significatively restored in the EditedΔ45–55 myotubes’ (46,5% of the nuclei) with higher values than control (37% of the nuclei) and ImΔ45**–**55-D1 myotubes’ (32% of the nuclei) (Fig. 5A,C).

Moreover, MyHC expression analysed by in-cell western observing a significant increase (near 2 times) in the edited differentiated myotubes’ compared to the DMD myotubes’ (Fig. 5D).

## DISCUSSION

The large in frame deletion including *DMD* exons 45 to 55 has been postulated as an attractive model for DMD therapy, because it could be beneficial to a high number of DMD patients harbouring different mutations and could advance asymptomatically or produce benign phenotypes. In addition, the resultant dystrophin maintains its filamentous structure and functionality despite the missing spectrin repeats in the central rod domain (Nicolas et al., 2015). However, this mutation is also associated with severe phenotypes, the responsible factors for which remain largely unravelled. Particularly, in this work we aimed to replicate, as a proof of concept, a del45**–**55 with specific intronic breakpoints (del45**–**55-D1) to recover dystrophin expression in a DMDΔ52 immortalized myoblasts using the CRISPR-Cas9 system. This specific mutation was shared by a significant set of patients (due to two founder events) with predominance of asymptomatic phenotypes (Poyatos-García et al., 2022). In addition, we obtained immortalized primary myoblasts from one of the asymptomatic patients sharing these specific breakpoints (ImΔ45**–**55-D1).

The del45**–**55-D1 consists on a 457.3 kb deletion, which preserves the promoters of the Dp140 and Dp116 dystrophin isoforms. Deletions disrupting the *Dp140* regulatory sequences in intron 44 have been associated to the risk of cognitive impairment and brain structural and functional abnormalities (Doorenweerd et al., 2014; Felisari et al., 2000) . In addition, this deletion conserved the expression of the lncRNAs, 44 and 44s2, reported as favourable by Gargaun and colleagues (Gargaun et al., 2021).

Although previous works have also used the CRISPR-Cas9 system to introduce the del45**–**55 (Ousterout et al., 2015; Young et al., 2016) they inserted larger deletions (688 kb and 725 kb, respectively) being able to target a greater number of patients, but these deletions are not based on natural models and probably, alter the regulatory elements located across introns 44 and 55 that might be relevant.

Even though the del45**–**55-D1 was successfully reproduced into a DMD-cell line and the edited clonal cell cultures were isolated, deep sequencing analysis revealed that the gRNA cleavage efficiencies were limited, as previously reported (Soblechero-Martín et al., 2021). This poor on-target efficiencies could have been caused by the myoblasts’ transfection difficulties (Ousterout *et al*., 2015; Wojtal *et al*., 2016), reflected by less than 15% of the DMD myoblasts transfected with the plasmids containing the CRISPR-Cas9 elements, expressed the GFP reporter. Alternatively, other strategies such as ribonucleoprotein (RNPs) complexes have demonstrated greater transfection and cleavage efficiencies as well as reduced off-target effects than plasmids (linked to the reduced time of active RNP complexes compared to plasmids) (Kim et al., 2014; Zhang et al., 2021). Indeed, our experience with RNPs in immortalized myoblast (but different loci) revealed cleavage efficiencies above 87% (Poyatos-García et al., 2023).

Despite the use of plasmid vectors to introduce the CRISPR-Cas9 system, no relevant off-target events were detected (Fig. S3, S4). However, deep sequencing analysis revealed the presence of substitutions distant to the expected Cas9 cleavage site, also in the non-transfected samples (Fig. S3). Interestingly, in all cases the substitutions were transitions (A>G or T>C), which correspond to the typical error of the Phusion™ High-Fidelity DNA Polymerase used in library generation. Besides, the frequency of the substitutions is below of the polymerase theorical error (Mcinerney et al., 2014), (Table S2). Thus, it is critical to perform deeper and unbiased analysis such as whole genome sequencing prior the application of this technology to the clinic.

Our experiments demonstrate that the generated EditedΔ45**–**55 clone harbouring the del45**–**55-D1 can restore the *DMD* reading frame and dystrophin expression. The production of large deletions might disturb the genomic architecture and alter the splicing process impacting on the resultant proteins and the phenotype (Tuffery-Giraud et al., 2017). We did not detect any alteration on the edited clone’s RNA, observing the expected transcript with exon 44 and 56 sequences juxtaposed (Fig. 3A).

On the other hand, the analysis of the dystrophin protein expression in the EditedΔ45**–**55 line was assessed by different techniques. First, by immunofluorescence we confirmed the dystrophin recovery and its correct location in the edited clones ‘differentiated myotubes. Dystrophin quantification was evaluated using in-cell western and western blot, confirming the dystrophin increase in the edited cells. In addition, for the western blot analysis we incorporated a total of 3 myoblasts from healthy male donors whose signal was grouped for the analysis to reduce variability, because as previously have been reported, we also observed variable dystrophin levels among the different healthy controls (Arechavala-Gomeza et al., 2009). Similar to the dystrophin cDNA quantification by ddPCR (Fig. 3B); here we observed that dystrophin protein levels in the EditedΔ45**–**55 cell line were higher than ImΔ45**–**55-D1 and even controls lines; confirming the variability of dystrophin expression across the different cell cultures. This variability is consistent with previous analysis performed in skeletal muscle sections from dystrophinopathic patients with different in-frame *DMD* mutations (Anthony et al., 2011b).

Finally, we confirmed that DMD myotubes present alteration of the differentiation process (evaluated through the calculation of the fusion index and the expression of MyHC), compared to control myotubes’. Both parameters were restored in the EditedΔ45**–**55 cell line harbouring the del45**–**55-D1.

Overall, although many uncertainties still exist surrounding the clinical variability of the *DMD* del45**–**55, we evaluated as proof of concept, the beneficial to mimic a specific intronic breaking points naturally present in a subgroup of subjects mostly running as benign. We have also recapitulated the ability of the versatile CRISPR-Cas9 system to create custom deletions, enabling the rescue of dystrophin expression as well as improving the altered phenotypes. In addition, the analysis was performed also comparing the edited cell line to the patient’s derived immortalized myoblasts with the same deletion, allowing the characterization and validation of both cellular models.

Thus, we consider that our replicative approach and our immortalized myoblast cell line may represent a valid model to undertake *in vitro* experiments, aimed to dilucidated unsolved DMD pathomechanisms.

## MATERIAL AND METHODS

### Cell cultures and myoblasts immortalization

HEK293 cells were maintained with Dulbecco’s modified Eagle Medium high glucose (DMEM), 10% Foetal Bovine Serum and 1% of penicillin-streptomycin (Thermo Fisher, Waltham MA, USA).

A myoblast immortalised DMD cell line with a deletion of exon 52 (DMDΔ52) (ID: DMD638a) was provided by the Institute of Myology (Paris, France) and the object of the CRISPR/Cas9 gene edition. Additionally, immortalised human myoblasts derived from three healthy male donors’ biopsies (ID: AB1079 (C1), AB1190 (C2) and AB678 (C3) of 38, 16 and 53 years old respectively) were also provided by the Institute of Myology.

To create an immortalised culture (named ImΔ45**–**55-D1), a biopsy of *tibialis anterior* from a 32-year-old male donor harbouring a del45**–**55 with specific intronic breakpoints (del45**–**55-D1), was obtained in La Fe University Hospital (Valencia, Spain) after informed consent; research ethics committee authorization 2018/0200). Primary human skeletal myoblasts were purified as previously described (Sabater-Arcis et al., 2021) and immortalized in collaboration with the Institute of Myology (Paris, France) to increase its proliferative capacity as follows: primary myoblasts were transduced with both hTERT and Cdk4 lentiviral vectors with a ratio of the number of transducing lentiviral particles to the number of cells (MOI) of 5 in the presence of 4 μg/ml of polybrene (Sigma-Aldrich, Sant Luis, MO, USA). Transduced cell cultures were selected with puromycin (0.2 μg/ml, Life Technologies, Carlsbad, CA, USA) for four days and neomycin (0.3 mg/ml, Life Technologies, Carlsbad, CA, USA) for ten days. Cells were seeded at clonal density and selected clones were isolated from each population using glass cylinders (Mamchaoui et al., 2011).

Human myoblasts were cultured with Skeletal Muscle Cell Growth Medium (SMC) (PELOBiotech, Planegg, Germany). Differentiation medium (DM), when needed, was prepared with DMEM, 2% of Horse Serum and 1% of penicillin-streptomycin (Thermo Fisher, Waltham MA, USA).

### Cell proliferation assay

A total of 5000 primary and immortalized myoblasts (derived from the patient 45**–**55) per well were seeded in 96-well plates and were incubated at 37 C in a humidified chamber with 5% CO_2_ for 24, 48, 72, and 96 h in SMC medium. Cell proliferation was measured using the CellTiter 96 aqueous non-radioactive cell proliferation assay (MTS) (Promega) as previously described (Poyatos-García et al., 2023).

### CRISPR-Cas9 design and gRNA selection

Two gRNAs targeting the vicinity of each breakpoint in intron 44 and 55 of the del45– 55-D1 (ChrX:32056814 and ChrX:31599476 respectively, according to the human genome reference GRCh37/Hg19), using the Zhang lab designing tool (crispr.mit.edu). The genomic location and the sequence of the four gRNAs can be found in Table S1.

Each gRNA was cloned into a plasmid containing spCas9 and EGFP sequences (PX458; Addgene 48138) (Ran et al., 2013). To test the cleavage efficiency of the sgRNAs, 1.5ug of each plasmid was transfected independently into HEK293 cells using lipofectamine 3000 (Invitrogen, Carlsbad, CA, USA). Forty-eight hours after transfection, the genomic DNA was extracted (QIAamp® DNA Mini Kit, Qiagen), amplified using primers hybridizing in the proximity of the cleavage site (Table S1), and the PCR products were purified. T7E1 assay was performed and indel frequencies were calculated as previously reported (Fuster-García et al., 2017).

The two gRNAs selected (gRNAs_44.1 and 55.2), targeting intron 44 and 55 breakpoints) were co-transfected into HEK293 cells (1.5ug of each plasmid) to evaluate the production of the *DMD* del45–55-D1. Forty-eight hours after co-transfection, cell’s DNA was extracted and amplified by PCR (del45–55-D1 screening primers) followed to Sanger sequencing (Table S1).

### Generation of edited clones with the del45–55-D1 (Edited**Δ**45–55)

Immortalized DMDΔ52 myoblasts were co-transfected with 1.5ug of each plasmid containing the selected sgRNAs using Viafect^TM^ (Promega) transfection reagent (1:5 ratio). Forty-eight hours after transfection, fluorescence activated cell sorter (FACS) was applied to seed individually the GFP positive cells into 96 well plates and amplified to form homogeneous clonal cell cultures as previously described (Soblechero-Martín et al., 2021). DNA was extracted from the successfully grown clones, analysed by PCR and resolved on a 1% agarose gel to detect those harbouring the del45–55-D1 (Table S1). One edited clone, named “EditedΔ45–55” was expanded for further for characterization analysis. These experiments were carried out at Biobizkaia HRI, Barakaldo, Spain (NAT-RD group).

### On-target and off-target analysis

For the selection of the potential off-targets that might be produced by the gRNAs_44.1 and 55.2 (containing up to three mismatches), we used the Cas-OFFinder tool (Bae et al., 2014).

Amplicon high-throughput sequence analysis was carried out to evaluate the on-target efficiencies and the potential off-target events. For the off-target analysis, we used DNA from DMD immortalized myoblasts co-transfected with both sgRNAs. For the on- target analysis DNA from DMD myoblasts transfected with the sgRNA of interest (either 44.1 or 55.2) was used in each case. DNA from DMD untransfected myoblasts was used as control for each target locus. Two-step PCR strategy was used to generate the library. For the first PCR (PCR1), primers for each locus contained an adapter sequence. PCR products were purified with AMPure Beads (BD Bioscience, US). For the second (PCR2), PCR products were re-amplified with primers containing the adapter sequence overlapping the first primers, and with an index sequence in the reverse primer. Final PCR products were purified with AMPure Beads (BD Bioscience). The library was prepared with PCR products pooled in equimolar amounts following the manufacture’s protocol and loaded in a Micro MiSeq Reagent Kit v2 (500-cycles) (Illumina, San Diego, CA, USA) on a MiSeq platform (Illumina). The fastaq.gz files were analysed through CRISPResso2 software to evaluate the edition efficiency and possible off target effects, using default parameters. The primers used for the PCR1 and PCR2 are listed in Table S3.

Additionally, we analysed the potential off-targets of the selected clone used for the functional characterization by Sanger sequencing (using PCR1 primers without the adapter sequence, Table S3).

### RNA analysis

RNA was extracted from differentiated cell cultures lines after being pelleted (RNeasy mini kit, Qiagen, Hilden, Germany). Reverse transcription was performed using 1 µg of total RNA and with SuperScript IV Reverse Transcriptase (Invitrogen, Waltham, MA), and nested PCR of cDNA samples was carried out using specific primer pairs (hybridizing in DMD exons 41 and 60 (RT-PCR1) and in exons 43 and 59 (RT-PCR2); Table S1) as previously described (Poyatos-García et al., 2022). PCR products were analysed on 1% agarose gels, DNA was purified (Gel Extraction Kit; Omega Bio-Tek, Norcross, GA) and validated via Sanger sequencing.

Duplex droplet digital PCR (ddPCR) was performed with 3µl of cDNA using the QX200 Droplet Digital PCR system (Bio-Rad Laboratories, Hercules, CA), as previously described (Poyatos-García et al., 2022), using the commercial probe dHsaCPE5049433, HEX labelled (Bio-Rad) for dystrophin quantification.

### Cell cultures Immunofluorescence

For all immunodetections, 2.5x10^4^ myoblasts /well were seeded in 24-well plates and, after seven days differentiating into myotubes, cells were fixed in 4% paraformaldehyde (PFA). Cell cultures were permeabilized with PBS-T (0.1% X-TritonX-100 in PBS 1X) and blocked for 1h at room temperature (RT) in PBS-T, 1% BSA, 1% normal goat serum (blocking buffer) before incubation with primary antibodies, diluted in blocking buffer, overnight at 4 °C. For dystrophin immunostaining, a mixture of three mouse monoclonal antibodies at 1:50 dilution was used: NCL-Dys1 (Novocastra Laboratories, Newcastle Upon Tyne, UK), Mandys1 and Mandys106 (The Wolfson Centre for Inherited Neuromuscular Disease). For myosin heavy chain detection (MyHC), a mouse monoclonal anti-MyHC antibody was used (MF20, 1:50, DSHB, University of Iowa, IA, USA) and for desmin detection, a rabbit polyclonal anti-desmin antibody (1:200, Abcam, Cambridge, UK). After washing with PBS-T, cells were incubated with the appropriate secondary antibody: goat anti-Mouse IgG (H+L) Alexa Fluor Plus 488 and goat anti-rabbit IgG (H+L) Alexa Fluor Plus 488 (1:200, Thermo Fisher). Finally, samples were mounted with VECTASHIELD® mounting medium containing DAPI (Vector Laboratories, London, UK) to detect the nuclei. Images were acquired in an LSM800 confocal microscope (Zeiss) at 200x magnification.

The fusion index was calculated as the percentage of nuclei within myotubes (>2 nuclei) out of the total number of nuclei in Desmin-positive cells (15 micrographs per condition). The differentiation index was calculated as the percentage of nuclei within MyHC-positive myotubes out of the total of nuclei (10 micrographs per condition).

### Protein quantification

In-cell western assays were performed as previously described (López-Martínez et al., 2022; Ruiz-Del-Yerro et al., 2018). Briefly, the cultures were seeded in 96-well plates and differentiated for 7 days. Plates then were fixed with ice-col methanol, permeabilised with PBS-T, blocked (blocking buffer, LI-COR® Biosciences) and incubated with primary antibodies overnight at 4°C. For dystrophin detection, the mixture of the three primary antibodies described above (NCL-Dys1, ManDys1 and ManDys106, 1:100 each) was used, and MF20 primary antibody (1:100) for MyHC detection. Next day, plates were incubated with the secondary antibodies. Biotin- mediated amplification (Abcam 6788 goat antimouse IgG biotin 1:2000) was used to increase dystrophin signal. Secondary antibodies, IRDye 800cw streptavidin (1:2000 and IRDye 800CW goat anti-mouse 1:500, were prepared together with CellTag 700 Stain (LI-COR® Biosciences) at 1:1000 dilution and incubated for 1 h at RT and protected from light. After incubation, plates were analysed using the Odyssey® CLx Imager (LI-COR® Biosciences).

For western blot quantification, cultures were seeded in P6 plates (2.5x10^5^ cells/well) and differentiated for seven days. Cell pellets were collected and solubilized in lysis buffer (Anthony et al., 2014b). Protein concentration was determined using the BCA Protein Assay (Thermo Fisher, Waltham MA, USA). Samples were loaded onto a NuPAGE® Novex® 3–8% Tris-Acetate (Thermo Fisher, Waltham MA, USA) and run at 100V during 5h. Protein wet transference onto 0.45 µm nitrocellulose membrane was carried out at 20V for 18h at 4 °C. Then, membranes were blocked in 5% non-fat dry milk diluted in TBST (0.1% Tween20) for 1.5 h at RT and incubated overnight at 4°C with primary antibodies: anti-dystrophin antibody (NCL-Dys1, 1:40, Novocastra Laboratories) and anti-alpha actinin antibody (A7732, 1:3000, Sigma-Aldrich). Membranes were incubated with the secondary antibody sheep anti-mouse IgG (H+L) (ab6808, 1:2000, Abcam) for 1h in dark. Membranes were revealed using SuperSignal™ West Pico PLUS (Thermo Fisher, Waltham MA, USA) using an Amersham Imager 600 (GE Healthcare, Chicago, IL) imaging system. Bands’ intensities were quantified with the ImageJ software (NIH, Bethesda, MD, USA). Dystrophin signal was normalized to alpha-actinin signal.

### Statistical analysis

Mann–Whitney U, Kruskal-Wallis and linear regression tests were used to determine the statistical significance of the obtained data. Statistical analysis was performed using GraphPad Prism 6 software.

## ACKNOWLEDGEMENTS

We strongly acknowledge the Fundación Isabel Gemio for supporting this work and for its enormous effort in securing funding, their trust in our work, and extraordinary cooperation.

We acknowledge the use of cell cultures provided by the Institut de Myologie (Paris, France). We gratefully acknowledge the use of the antibodies provided by Professor Glenn Morris from the Muscular Dystrophy Association (MDA) Monoclonal Antibody Resource, which distributes antibodies for research in neuromuscular diseases worldwide from Oswestry, United Kingdom. The MF20 antibody developed by D.A. Fischman, Weill Cornell Medical College, was obtained from the Developmental Studies Hybridoma Bank, created by the NICHD of the NIH and maintained at the University of Iowa, Department of Biology, Iowa City, IA 52242. Cell sorting experiments were performed at the Cell Analytics Facility at Achucarro-Basque Centre of Neuroscience (Leioa, Spain).

## DATA AVAILABILITY

All relevant data can be found within the article and its supplementary information. All materials and further information of this study is available upon request.

## AUTRHOR CONTRIBUTION STATEMENT

Conceptualization, J.J.V., J.P.-G., R.P.V.-M., V.A.-G.; software, A.L, E.G.-R., G.G.-G; methodology, validation, formal analysis and investigation, J.P.-G., P.S.-M., A.L., A. L.- M.; E.G.-R., G.G.-G.; resources, A.L., R.P.V.-M., N.M, G.G.-G., J.O., V.A.-G.; writing-original draft preparation, J.P.-G., J.J.V.; writing-review and editing, J.P.-G., P.S.-M., A.L., A. L.-M.; E.G.-R., R.P.V.-M., N.M., G.G.-G, J.O., V.A.-G., J.J.V.; visualization, J.P.-G, P.S.-M., E.G.-R.; supervision, J.J.V., A.L., R.P.V.-M., V.A.-G; project administration, J.J.V, V.A.-G. and funding acquisition, J.J.V., V.A.-G.

